# Computational hypothesis: How Intra-hepatic functional heterogeneity may influence the cascading progression of free fatty acid-induced non-alcoholic fatty liver disease (NAFLD)

**DOI:** 10.1101/2020.05.28.120626

**Authors:** Hermann-Georg Holzhütter, Nikolaus Berndt

## Abstract

Non-Alcoholic Fatty Liver Disease (NAFLD) is the most common type of chronic liver disease in developed nations, affecting around 25% of the population. Elucidating the factors causing NAFLD in individual patients to progress in different rates and to different degrees of severity, is a matter of active medical research. Here, we aim to provide evidence that the intra-hepatic heterogeneity of rheological, metabolic and tissue-regenerating capacities plays a central role in disease progression. We developed a generic mathematical model that constitutes the liver as ensemble of small liver units differing in their capacities to metabolize potentially cytotoxic free fatty acids (FFAs) and to repair FFA-induced cell damage. Transition from simple steatosis to more severe forms of NAFLD is described as self-amplifying process of cascading liver failure, which to stop depends essentially on the distribution of functional capacities across the liver. Model simulations provided the following insights: (1) A persistently high plasma level of FFAs is sufficient to drive the liver through different stages of NAFLD; (2) Presence of NAFLD amplifies the deleterious impact of additional tissue-damaging hits; and (3) Coexistence of non-steatotic and highly steatotic regions is indicative for the later occurrence of severe NAFLD stages.

## Introduction

Non-Alcoholic Fatty Liver Disease (NAFLD) is the most common type of chronic liver disease in developed nations, where it affects around 25% of the population [1]. NAFLD starts with simple steatosis and may progress to severe liver diseases like cirrhosis or hepatocellular carcinoma (HCC). Up to 30% of patients with liver steatosis develop a Non-Alcoholic Steatohepatitis (NASH), an inflammatory state of the liver that in about 20% of cases develops further to cirrhosis and end-stage liver failure [1]. Owing to the high metabolic reserve capacity of the liver, progression of NAFLD typically proceeds silently over long periods before an abrupt worsening of central metabolic liver functions results in severe clinical symptoms requiring urgent treatment.

Steatosis of the liver in the absence of significant alcohol intake is commonly associated with insulin resistance resulting from hyper-caloric diet and a sedentary lifestyle. Insulin resistance of the adipose tissue increases the release of free fatty acids (FFAs) into the blood [2] and thus promotes the uptake of FFAs into liver, kidney, heart and other organs using FFAs as preferred energy-delivering substrates. FFAs exceeding the cellular need for ATP synthesis are esterified to triacylglycerol (TAG) and membrane lipids. Conversion of excess fatty acids into complex lipids serves as a detoxification mechanism [3-6] as elevated levels of FFAs and some of their reaction products may cause cell damage [7]. As shown in mice with downregulated diacyglycerol acyl transferase 2 (DGAT2), a rate-limiting enzyme of TAG synthesis, inhibition of TAG synthesis improved hepatic steatosis but exacerbated liver damage exerted by elevated levels of potentially cytotoxic FFAs [6]. FFA-induced hepatotoxicity includes various molecular mechanisms as, for example, induction of endoplasmic reticulum stress, generation of free radicals and subsequent activation of the mitochondrial apoptosis pathway [8]. Necrosis and apoptosis of hepatocytes mount a dynamic multicellular response wherein stromal cells are activated *in situ* and/or recruited from the bloodstream, the extracellular matrix is remodeled, and epithelial cells expand to replenish their lost numbers [9]. If the FFA challenge persists, a quasi-stationary balance between damage and tissue regeneration is established, which over the time may slowly shift towards fewer and fewer vital hepatocytes and more and more non-functional fibrotic lesions [9].

Several factors may explain why patients with diagnosed steatosis may or may not develop NASH and even more severe chronic liver disease. As with all diseases, genetic factors may determine the susceptibility of an organ to damaging events. Genome-wide association studies have revealed several single nucleotide polymorphisms associated with the pathology of NAFLD, among them the gene variant I148M of the enzyme Patatin-like phospholipase domain-containing 3 playing an important role in the cellular lipid and lipid droplet metabolism [10]. On top, the liver is continuously confronted with all kinds of orally administered toxins and gut-derived pathogens.

A now widely accepted hypothesis postulates that simple steatosis (‘first hit’) has to be followed by further pathophysiological hits, such as pathogen-associated acute inflammation or gut-derived endotoxins, to push the liver successively into a critical state [11]. A third factor influencing the progression of NAFLD consists in the intra-hepatic spatial heterogeneity of metabolic and immune-modulatory capacities. Gradients along the porto-central axis of the liver lobule exist not only for genes and proteins of hepatocytes leading to zone dependent differences in lipid metabolism [12] but also in other types of liver cells and the matrix of the space of Disse [13, 14]. Zone-dependent differences in gene expression within a single lobule can be partially accounted for by concentrations gradients of metabolites, hormones and morphogens along the sinusoidal blood stream [15, 16]. In adults, steatosis is most intense around the central veins (predominantly in zones 2 and 3) whereas children may have an alternate pattern of progressive NAFLD characterized by a zone 1 distribution of steatosis, inflammation and fibrosis [17].

Besides intra-lobular metabolic heterogeneity, independent experimental findings point to another layer of intra-hepatic heterogeneity that hitherto has not been implicated in NAFLD progression. We will refer to this type of heterogeneity as ‘macro-scale heterogeneity’ as it runs on lengths scales of millimeters and centimeters, which are orders of magnitude above the size of a single liver lobule. Support for the existence of macro-scale heterogeneity comes mainly from functional liver imaging and histological findings. High-resolution imaging of hepatic blood flow revealed large regional variability up to a factor of three [18, 19]. Hepatic clearance of 2-[18^F^]fluoro-2-deoxy-D-galactose measured by PET/CT varied by about 24% in patients and 14% in healthy subject [20]. The spatial distribution of liver fat in adults with NAFLD showed a variability in the range of 0.7-4.5% [21]. Patient-specific spatial pattern of fat dispositions may largely vary from diffuse fat accumulation with and without focal sparing and focal fat accumulation [22]. Presence of macro-scale heterogeneity in functional and metabolic parameters is also reflected by differences of NAFLD-associated histopathologic lesions between different anatomical parts of the liver [23].

Taken together, progression of NAFLD can be affected by several intrinsic and environmental factors, among which the rise of the serum FFA level, the distribution of intra-hepatic functional capacities, and the eventual occurrence of additional liver-damaging events play a central role. The question of how these factors may act together in the patient-specific development of NAFLD can hardly be addressed in a clinical approach for obvious technical and ethical reasons. Computer simulations based on physiologically reliable mathematical models may serve as an alternative in such situation. This prompted us to develop a generic and in many ways further expandable mathematical model of NAFLD progression.

## Materials & Methods – Mathematical Model

The mathematical model used for our disease simulations consists of three modules. (1) The hemodynamic module describes the blood flow within the liver sinusoids as was previously described [24]. (2) The metabolic module describes the cellular turnover of FFAs and TAG, and (3) the damage-repair module describes the FFA-induced damage of hepatocytes and tissue repair.

### Module 2: Kinetic model of FFA and TAG turnover in a liver unit

We subdivided the liver into a larger number of small liver units (LUs) differing in their hemodynamic properties, metabolic capacities, vulnerability against FFA-induced cell damage and wound-healing capacities. A single LU is described by a two-compartment model: The vascular compartment formed by the sinusoids receives fatty acids with the blood from the hepatic arteries and exchanges fatty acids with the cellular compartment, which is equipped with the average metabolic capacities of all hepatocytes in the unit (see Fig. 1). The numbers on the reaction arrows in Fig. 1B are FFA particle numbers (in micromoles) and particle transport rates (in micromoles/100 ml/min) for a healthy LU with a volume of 100 ml. They represent typical values of transport and metabolization rates of FFAs measured in the healthy human liver (see references in Table 1)

**Table 1.** Rate laws for lipid fluxes and values of kinetic parameters for a standard liver unit. Rate constants were based on the particle numbers and particle transport rates shown in Fig. 1. *) calculated from the published VLDL-TAG secretion rate [25] by assuming a liver volume of 1350 ml.

| Flux | Rate law | Model value<br>($\mu\text{mol}/\text{min}/100 \text{ ml}$) | Experimental value<br>($\mu\text{mol}/\text{min}/100 \text{ ml}$) | Rate constant<br>($\text{min}^{-1}$) |
| --- | --- | --- | --- | --- |
| $v_u$ | $v_u^i = k_u \cdot (fa_{s^i} - fa_i)$ | 7 | 10 [26]<br>0-40 [27] | $k_u = 0.78$ |
| $v_{denovo}$ | $v_{denovo} = k_{denovo}$ | 1 | 3%-18% $v_u$ [28] | $k_{denovo} = 0.012$ |
| $v_\beta$ | $v_\beta = k_\beta \cdot fa_{c^i}$ | 4.5 | 5 [29] | $k_\beta = 0.6$ |
| $v_{+tag}$ | $v_{+tag} = k_{+tag} \cdot \left(\frac{fa_{c^i}}{fa_{crit}}\right)^2$ | 14 | 2-6 [30]<br>0-20 [31] | $k_{+tag} = 3.6$ |
| $v_{-tag}$ | $v_{-tag} = k_{-tag} \cdot tag^i$ | 11 | | $k_{-tag} = 0.006$ |
| $v_{VLDL}$ | $v_{VLDL} = k_{VLDL} \cdot tag^i$ | 3.5 | 4.5 *) [32]<br>1.5-10 [33] | $k_{VLDL} = 0.018$ |

**Fig 1.**
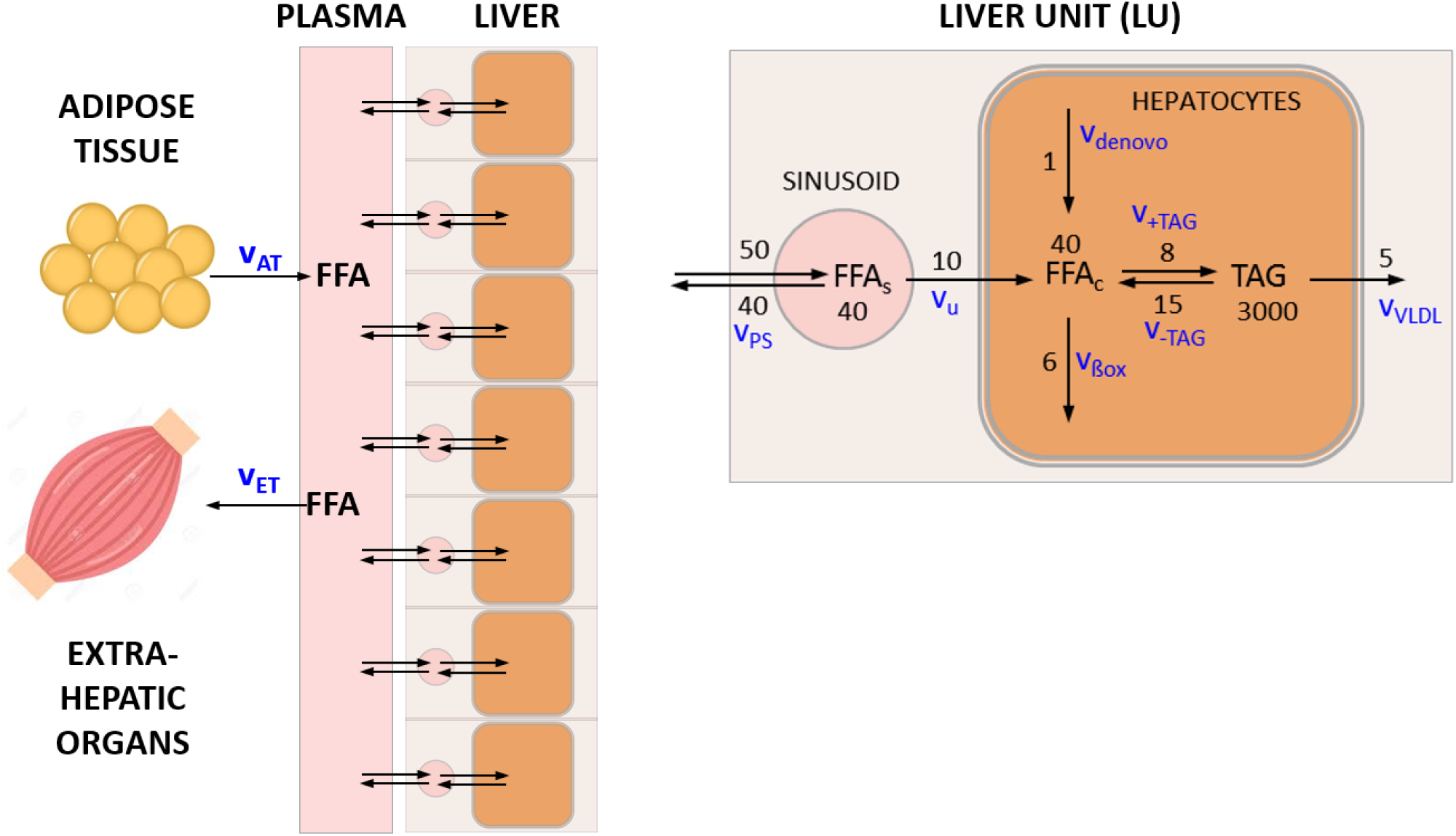
Transport and metabolism of free fatty acids (FFAs) in the liver. A) The liver is composed of a given number (N_U_) of liver units (LUs), which differ in their functional capacities. Each LU comprises a vascular compartment and a cellular compartment. Plasma fatty acids (FFA_p_) result mainly from lipolysis in the adipose tissue (release rate v_AT_). They can be utilized by various extra-hepatic organs, in particular the skeletal muscle (utilization rate v_ET_). B) FFAs enter the vascular bed of the liver with the rate v_ps_ that is determined by the plasma concentration of fatty acids and the blood flow rate. A certain fraction of fatty acids is taken up by hepatocytes with rate v_u_, the remaining part re-enters the circulation via the venous blood efflux. Cellular fatty acids (FFA_c_) are either formed *de novo* (synthesis rate v_denovo_) or taken up from the plasma. They can be used as building blocks for the synthesis of various lipids, mainly triacylglycerols (TAG) (synthesis rate v_+TAG_). Note that the variable TAG denotes the pool of FFAs esterified in TAG in the model. Alternatively, FFA_c_ can be degraded to acetyl-CoA, either by mitochondrial and peroxisomal β-oxidation or CYP-450-mediated ω-oxidation (oxidation rate v_ßox_). TAG, the most abundant cellular lipid, is stored in lipid droplets, which can by hydrolyzed to FFA_c_ if needed (degradation rate v_-TAG_). A fraction of TAG is loaded in VLDL lipoproteins and exported (export rate v_VLDL_). The numbers on the reaction arrows are FFA particle numbers (in micromoles) and particle transport rates (in micromoles/100 ml/min) for a healthy LU with a volume of 100 ml. The corresponding mass transport rates are given in Table 1.

The kinetics of FFAs in the plasma and in the i-th LU (i=1,…,N_LU)_ is governed by the following system of first-order differential equations:

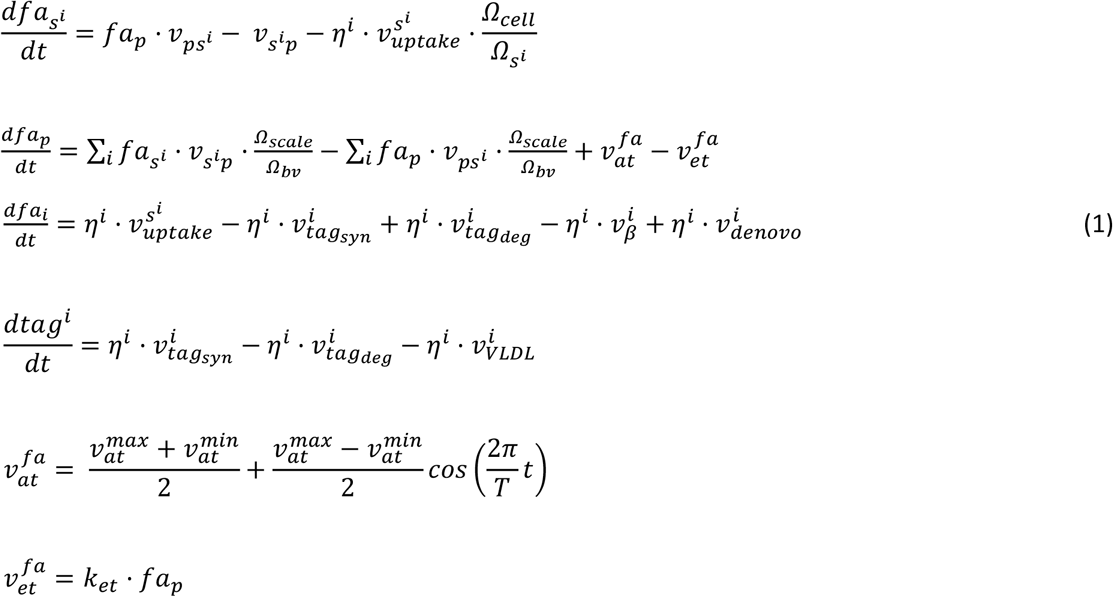

The blood flow is given by:

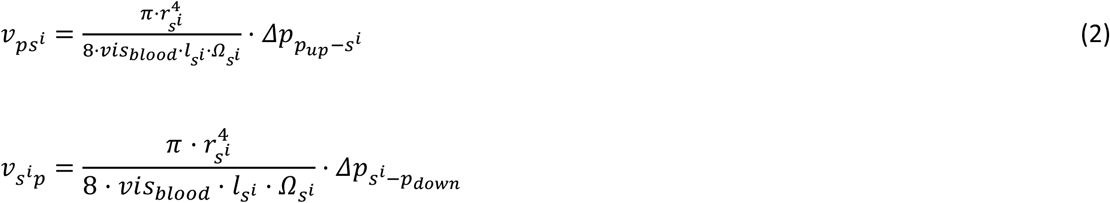

The upper index (i) numbers the LUs 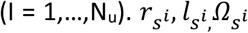 denote the radius, length and volume of the i-th LU, *vis*_*blood*_ is the blood viscosity, 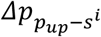 is the pressure gradient between arterial blood and the i-th LU and 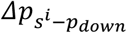 denotes the pressure gradient between venous blood and the i-th LU. For details, see Berndt et al., 2018 [24].

The parameter η denotes the fraction of metabolically intact hepatocytes in the LU. The quadratic term on the right-hand side of the kinetic equations for TAG synthesis takes into account that with increasing cellular content of FFAs, the synthesis of TAG and storage in lipid droplets proceeds super-linear due to the upregulation of lipogenic enzymes [34]. *fa*_*crit*_ denotes the critical cellular concentration of fatty acids, at which the rate of TAG synthesis starts to rise in a non-linear fashion. Note that the time-dependent variation of plasma FFAs (*fa*_*p*_) depends on the contribution of all LUs to the elimination of FFAs from the plasma.

Fig. 2 illustrates how differences in the metabolic capacities of individual LUs may influence the diurnal level of cellular FFAs. The rate of fatty acid influx into the plasma was modeled by a periodic function with period T, reflecting the insulin-dependent variation of FFA release from the adipose tissue to the plasma.

**Fig 2.**
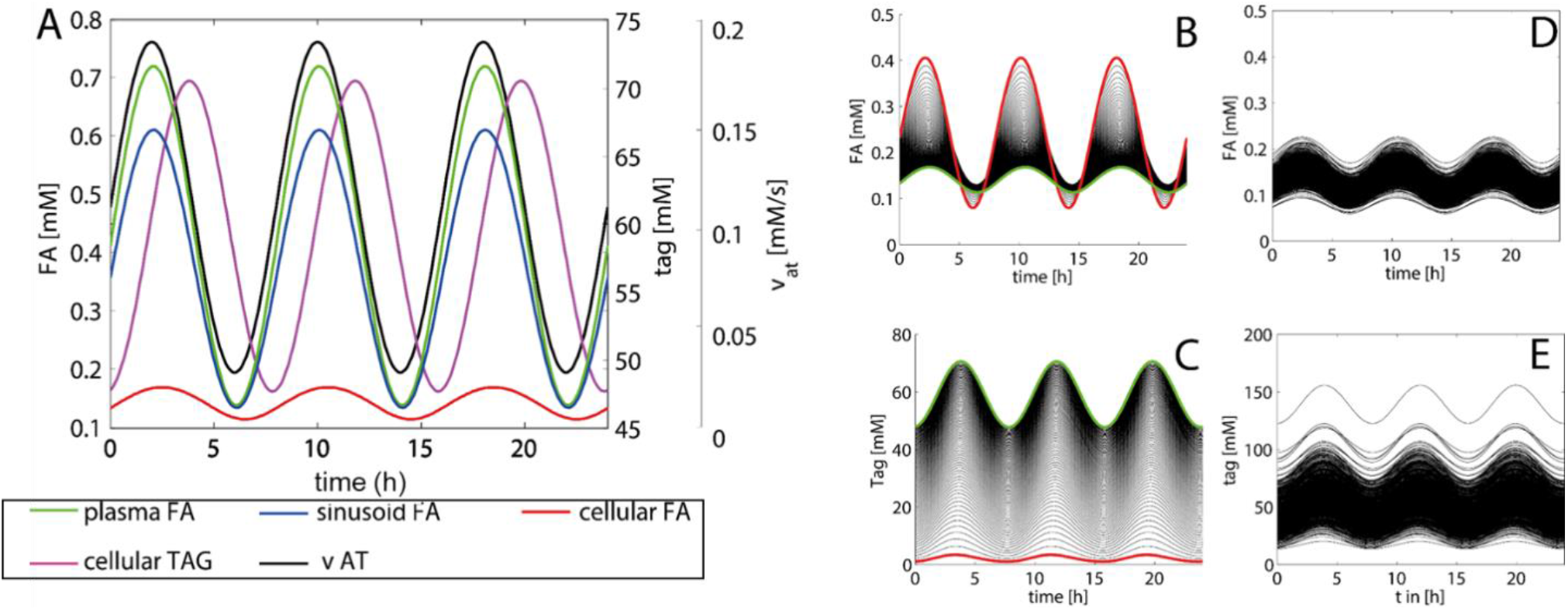
Diurnal variations of the concentration levels of FFAs and TAG. In this example, the release rate of FFA to the plasma varies three times per day between 1.8 mM/min and 11.4 mM/min resulting in a variation of the plasma concentrations of FFAs within the physiological range [0.4 mM; 0.7 mM] [35]. A) Time-course of FFAs and TAG in a standard LU endowed with the kinetic parameters given in Table 1. B,C) Impact of the cellular TAG storage capacity on variations of cellular TAG and FFAs. The curves were generated by varying the rate constant k_+TAG_ for TAG synthesis in small steps between 0.036 min^-1^ and 3.6 min^-1^. The values of all other model parameters were the same as given in Table 1. Red curves: k_+TAG_ = 0 (no TAG synthesis). Green curves: k_+TAG_ = 3.6 min^-1^ (highest TAG synthesis) C,D) Time-course of FFAs and TAG at randomly varied kinetic parameters. 100 curves were generated with random, normally distributed parameter values having as sample mean the original values in Table 1 and possessing a standard deviation of 20% (= 0.2 mean value).

The amplitude of variations of the lipid species decreased in the order FA_p_ *→* FA_s_ *→* FA_c_ *→* TAG. Figs. 3B,C illustrate how daily concentration changes of cellular FFAs depend on the TAG-synthesizing capacity. Increasing the cellular capacity to convert FFAs into TAG reduces the amplitude of diurnal variations and the 24h mean of the cellular FFA concentration, but increases the amplitude of variations and the mean of the TAG content. Note that at normal rate of TAG synthesis the daily maximum of cellular TAG is about 1.8 fold higher than the minimum (cf. green curve in Fig. 2C). This difference is in good agreement with measured TAG variations between the fasted and the fed state [36]. Figs. 3D,E illustrate the effect of random variations of metabolic parameters on the cellular profile of FFAs and TAG. 20% of random variations of the reference parameters (given in Table 1) may give rise to concentration maxima of cellular FFAs between 0.08 mM and 0.68 mM. For the variation of TAG, the coefficient of variation (CV; standard deviation to mean) was 35%, which is in good concordance with values obtained in outbred liver-healthy Wistar rats [37].

**Fig 3.**
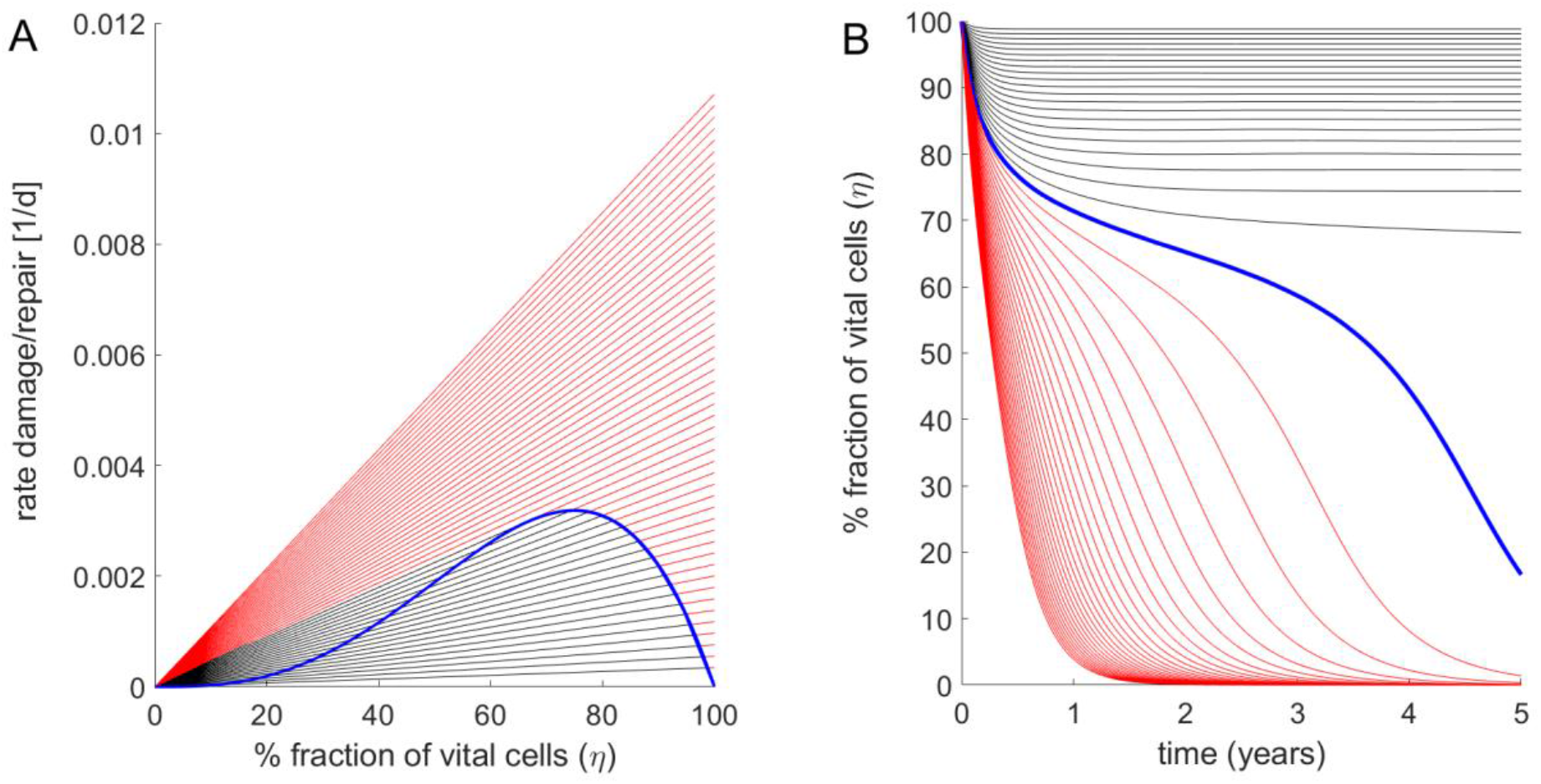
Fraction of intact hepatocytes in a single LU at increasing strengths of the FFA challenge. A) Rate of cell regeneration (blue curve) rates of cell damage at various FFA concentrations (straight lines) of a LU versus fraction of vital intact cells. Parameter values: *fa*_*crit*_ = 0.1 *mM*; k_d_ = 3.456·10^−4^ d^-1^; k_r_ = 3·10^−2^ d^-1^; β =30. Starting with 100% intact cells, the regeneration rate v_r_ increases with decreasing cell number to reach a maximum for η = 0.66. Below this critical value, the regeneration rate decreases. Straight lines refer to damage rates at varying concentrations of cellular FFAs. The ratio 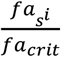 between the cellular FFA concentration and the critical concentration above which cell damage occurs was increased from 1 to 3 in steps of 0.1. The black lines hold for ratios 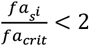 where stable stationary states with non-zero fractions of intact cells exist (given by the intercept points between the black lines and the ascending part of the blue curve). For ratios 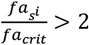 (red lines) the FFA-induced damage rates are persistently larger than the regeneration rate (no interception points between the red lines and the blue curve exist), i.e. the stationary state is reached at complete depletion of intact cells (η = 0). B) Time-dependent changes of the fraction of vital cells η if an initially fully intact liver unit (η = 1) is challenged with an elevated concentration of cellular fatty acids. As in A), the ratio 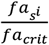 was increased from 1 to 3 in steps of 0.1. With increasing time, the time course of the vital cell fraction converges towards to the stationary value defined by the steady-state lines in A).

### Module 3: Fatty-acid induced cell damage and tissue repair

Numerous studies have provided evidence that elevated levels of fatty acids and of lipid species derived from fatty acids may cause cell damage resulting ultimately in cell loss by necrosis or apoptosis [38, 39]. Enhanced damage and loss of cells elicits a regenerative response, which in the liver includes mitotic division of hepatocytes and in case of severe damage also the differentiation of hepatic stem cells. In the model, time-dependent changes of functionally intact cells in the i-th LU was described by the following equations:

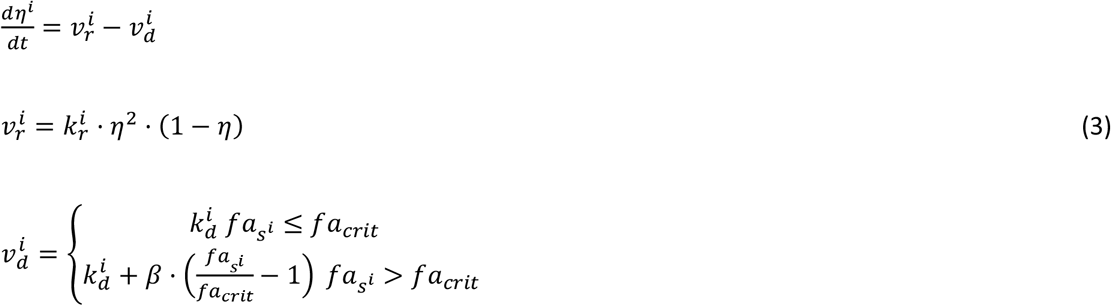

Here, η _i_ is the fraction of functionally intact hepatocytes, 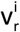 and 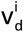denote the rates of cell regeneration and cell loss of the i-th LU. v_r_ was chosen to capture two opposing effects: The factor (1 − η) takes into account that the regenerative response initiated by immune cells rises with increasing cell damage, the factor η^2^ takes into account that tissue regeneration requires interacting cells. The combination of the two effects entails that the rate of tissue regeneration v_r_ is a non-monotone function with respect to η (represented by the green curve in Fig. 3A). The rate v_d_ of cell damage follows a first-order kinetics given by the cellular damage rate FD times η (represented by the black and red lines in Fig. 3A). FD is the sum of a basal rate k_d_ accounting for the continuous but small cell loss in the healthy liver, and an additional FFA-dependent term. The parameter β determines the increase of v_d_ if 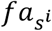 exceeds the threshold *fa*_*crit*_.

The steady-state fraction of active hepatocytes is given by the condition v_r_ = v_d_ which in Fig. 3A corresponds to the intersection points between the green curve and the straight lines. If the damage rate exceeds a critical value, the fraction of active cells tends to zero (see Fig. 3B). The numerical values of *fa*_*crit*_ and of the kinetic parameters k_d_, k_r_ and β (see legend of Fig. 3) were chosen such that for the healthy liver, the life-span of hepatocytes amounted to 365 days (1 year), whereas for the cirrhotic liver and a fatty acid concentration that is twice as high as the critical concentration *fa*_*crit*_, i.e. FFA_c_ = 2. *fa*_*crit*_, the life span is shortened to 26 days according to findings in Macdonald, 1961 [40].

## Results

### Simulation of FFA-induced NAFLD progression

Simulations of FFA-induced NAFLD progression of the whole liver were performed by numerical solution of the coupled equations (1) and (2) for each LU. As the progression of NAFLD takes place on the time scale of months and years, in these simulations the daily variations of lipid species (see Fig. 2) were neglected and their concentration values were put to the quasi-stationary 24h time averages. Functional heterogeneity of the liver was taken into account by randomly varying the kinetic parameters of the LUs by 20%, which corresponds to typical variations of blood flow [41] and tissue variations of protein abundances [42, 43]. The simulations started at t = 0 with a healthy liver and a physiologically normal FFA release rate of 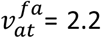 mM/min to the plasma. At time t = 5 years, the release rate was increased to 3.3 mM/min. This resulted in a rise of plasma FFAs and the onset of liver steatosis, which was quantified as percentage of LUs having a TAG content larger than 30 mM. In concordance with lipidomics data of human NAFLD and NASH livers [44], the simulations yielded an about 6-fold initial rise of steatosis followed by a moderate decline with progressing NAFLD.

Severity of NAFLD at time t was evaluated by the total fraction of intact hepatocytes (TFH),

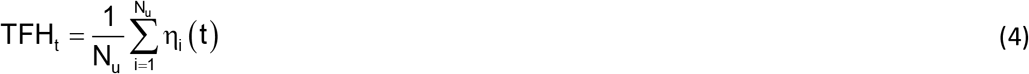

At early time points after onset of the FFA challenge, LUs endowed with the lowest capacities for TAG storage and/or tissue repair are unable to establish a new balance between damage and repair and thus lose their functionality, i.e. η tends to zero. The metabolic failure of these units entails a reduced overall FFA-esterifying capacity of the liver, which causes an additional rise of the FFA plasma level and thus increases the FFA burden to the remaining intact LUs. This self-amplifying cycle of organ damage may stop if a remaining fraction of intact LUs exists that are endowed with sufficient metabolic and repair capacities to cope with the elevated levels of FFAs.

Although the magnitude of the random variations of model parameters was identical (standard deviation = 20%), the severity of NAFLD after 30 years displayed large differences (see Fig. 4C). The reason for this heterogeneity is that the parameters determining blood flow rate, uptake and removal of FFA (k_u_, k_ß_, k_+TAG_, k_-TAG_) and damage and repair (k_d_, k_r_) vary independently from one LU to another. Hence, the number of LUs endowed with the most favorable combination of these parameters (fast blood flow + high rate of FFA removal + low damage rate + high repair capacity) may differ from liver to liver.

**Fig 4.**
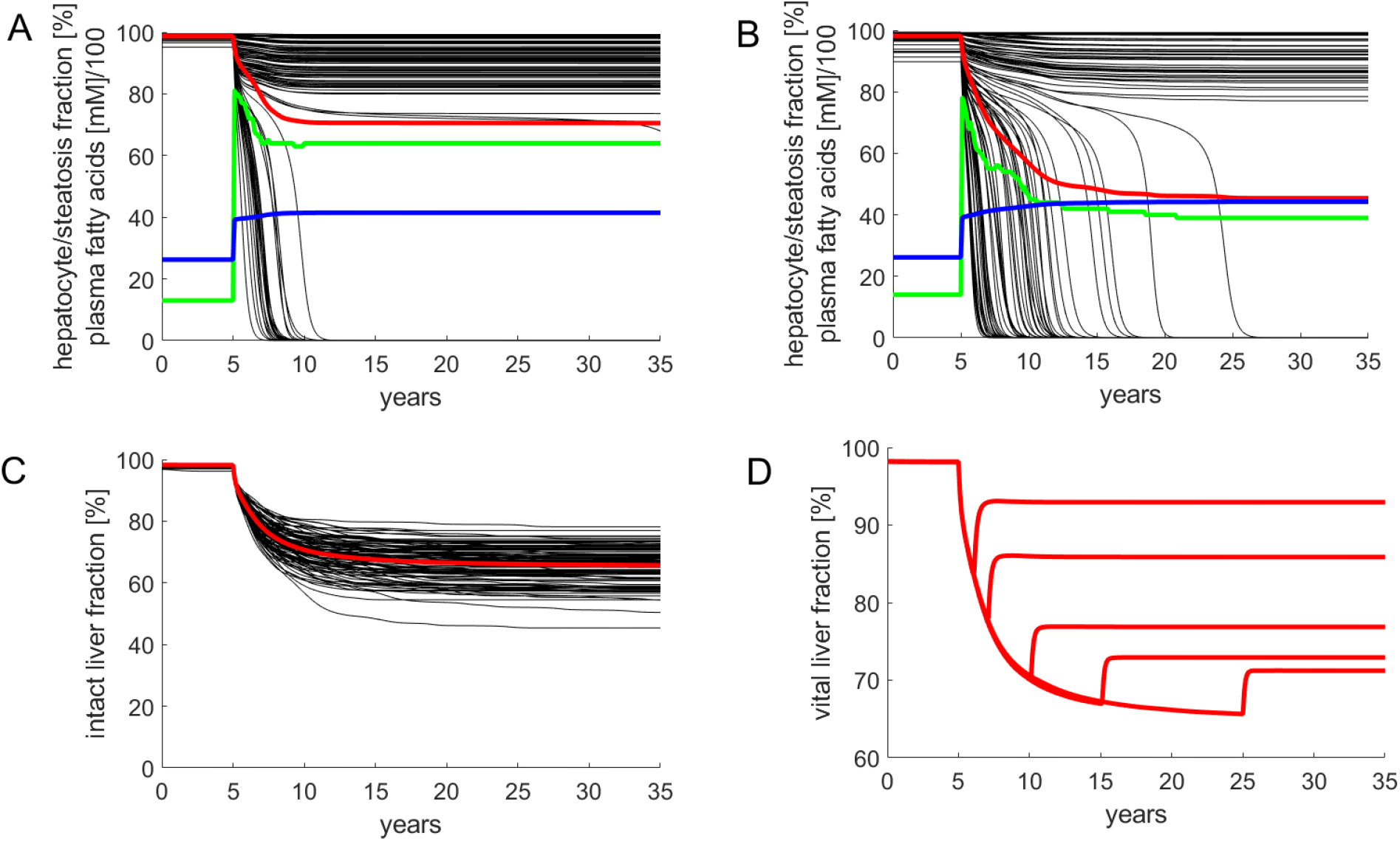
Simulated progression of FFA-induced NAFLD. Number of liver units N_u_ = 100; Volume of a single LU = 15 ml. The kinetic parameters of individual LUs varied randomly with a standard deviation of 20% around the reference values given in Table 1. The release rate of FFA into the plasma was put to the reference value of 2.2 mM/min corresponding to a plasma FFA concentration of 0.26 mM. The simulation of the healthy liver was run until the concentration of all model variables (concentration of lipid species and fractions of active hepatocytes) had reached stationary values. At time t = 5 years, the liver was confronted with a plasma FFA challenge elicited by an increase of the FFA release rate to 3.3 mM/min. This resulted in an initial increase of the FFA plasma concentration to 0.42 mM. Note that the further increase of the FFA plasma concentration is due to the progressive decline of the total fraction of intact hepatocytes and the concomitant decline of the total hepatic FFA utilizing capacity. A,B) Examples for the variable response of the liver to an identical FFA challenge. Black curves; Fraction of intact hepatocytes in individual LUs. Red curves: Total fraction of intact hepatocytes (TFH) defined by equation (4). Blue curves: FFA plasma concentration (FFA_p_). Green curves: Liver steatosis defined through the fraction of liver units having a mean TAG content larger than 30 mM. C) Progression of NAFLD in 100 model livers. D) Simulated residual regenerative capacity of the liver when the FFA challenge was ceased at the indicated time points after onset of NAFLD

For the example in Fig. 4A, the overall fraction of active hepatocytes dropped within five years to a moderately reduced and stable level of 80%. The clinical correlate of this type of NAFLD is a steatotic liver with low-grade inflammation. For the example in Fig. 4B, the overall fraction of active hepatocytes dropped continuously to 45%. Clinically, this time course corresponds to disease progression through the stages steatosis *→* NASH *→* cirrhosis.

The example in Fig. 4D illustrates how the liver may recover after cessation of the FFA challenge. Without lowering the release of FFAs into the plasma, the THF declines within 30 years to about 65%. This can be prevented by interrupting the FFA challenge, i.e. setting the release rate of FFA into the plasma back to the normal value (2.2 mM/min in the model). A short FFA challenge of several month, as it may occur during total parenteral nutrition [45] or transient insulin resistance due to infections [46] had virtually no impact on the hepatocyte fraction. The longer the FFA challenge persisted, the lower was the fraction of hepatocytes, which could be rescued from further damage. After five years of ongoing NAFLD, interruption of the FFA challenge restored only 77% of the functional intact liver mass. The time courses in Fig. 4D are in good concordance with reported findings according to which the regeneration of fibrotic NAFLD livers after removal of the lipid load was generally incomplete and took many months [47, 48].

In all simulations, steatosis was generally highest immediately after onset of the FFA challenge. The decrease of hepatic TAG with progressing NAFLD reflects the decrease of active cells capable of synthesizing and storing TAG. The more aggressive NAFLD, the faster was the decline of steatosis.

### Size and length scale of intra-hepatic parameter heterogeneity influences NAFLD progression

The simulations of NAFLD progression shown in Fig. 4 were performed for livers composed of 100 LUs. With a liver volume of 1500 ml, this means a LU volume of 15 ml, which corresponds to a spherical LU with a diameter of about 3 cm. This is a typical length scale, at which significant spatial differences in liver steatosis are commonly assessed by means of MRI techniques [49, 50]. In order to check how strong the statistical differences between individual NAFLD time courses depend on the size and length scale of parameter heterogeneity, we carried out simulations with livers composed of 10, 100 and 1000 LUs, corresponding to a characteristic length scale of 6.6 cm, 3 cm and 1.4 cm, respectively. As before, random parameter variances were put to 20%. As endpoint, we used TFH_30_, the total fraction of intact hepatocytes at 30 years after onset of the FFA challenge. For comparison, we carried out the simulations also for completely homogenous livers composed of LUs with identical but from liver to liver randomly varying parameter sets. Intriguingly, the homogeneous liver showed an ‘all-or-none’ characteristics of NAFLD progression: 90% of livers had none or only marginal signs of NAFLD but the remaining 10% ended up in complete liver failure (Fig. 5A). In contrast, complete liver failure was not observed for the heterogeneous livers also composed of 100 LUs but the distribution of TFH_30_ values was much broader compared to the homogenous liver (Fig. 5C). Hence, intra-hepatic functional heterogeneity prevents complete liver failure but entails on the average a more severe NAFLD progression as compared with a functionally homogeneous liver. Increasing the number of LUs, the distribution of THF_30_ values became narrower and the mean THF_30_ values were getting smaller than the THF_30_ of the homogeneous liver (indicated by the red bars in Fig. 5) From this can be concluded that the risk for NAFLD progression to severe forms increases with an increasing length scale of functional heterogeneity.

**Fig 5.**
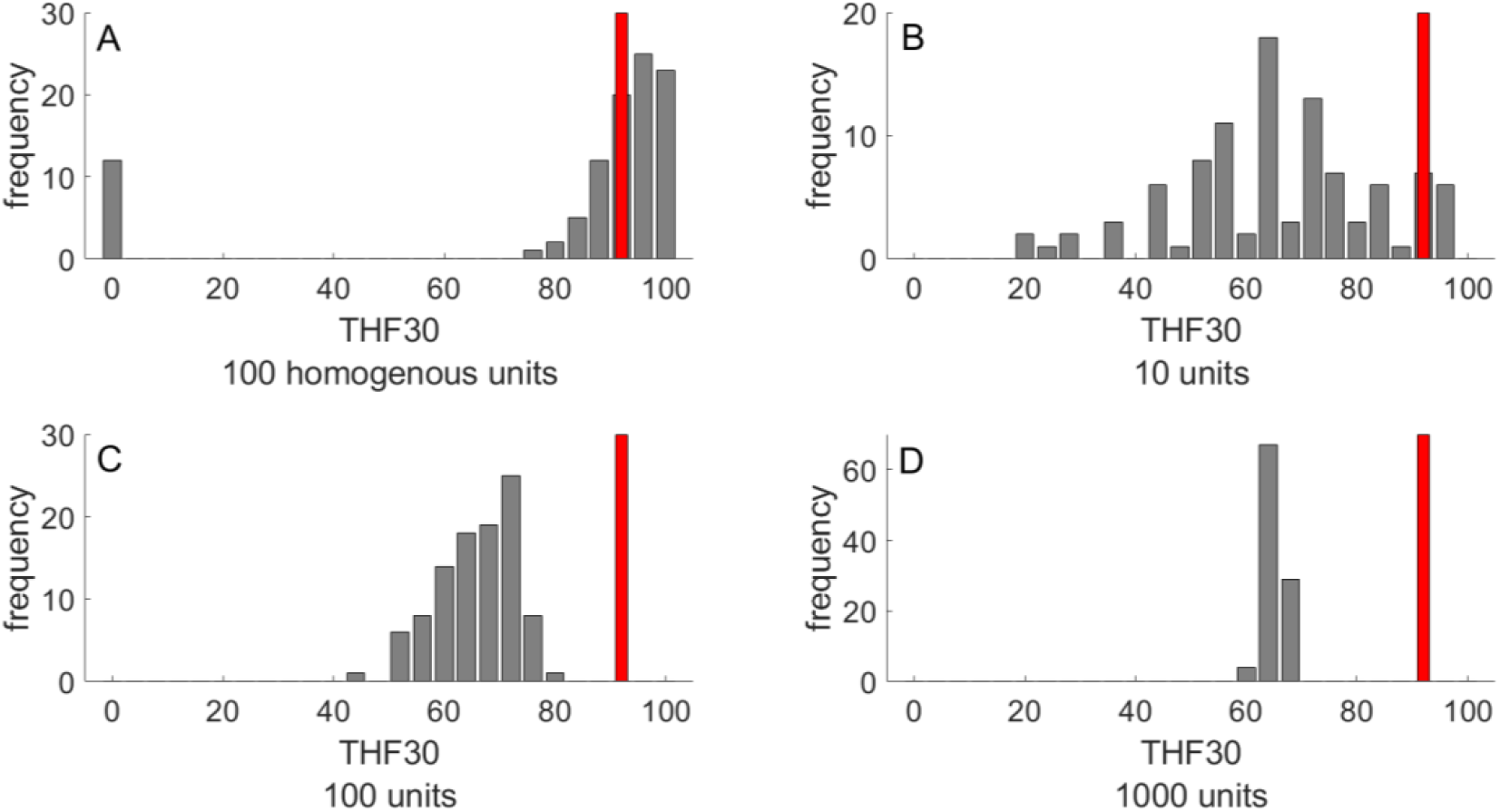
Outcome of NAFLD progression in 100 livers after 30 years NAFLD progression The outcome was evaluated in terms of TFH_30_ representing the total fraction of vital hepatocytes after 30 years of FFA challenge. A) Homogenous liver composed of N = 100 functionally identical LUs. B-D) Heterogeneous livers composed of various numbers (10, 100, 1000) of functionally LUs. The parameter set for each LU was randomly chosen from a normal distribution with standard deviation SD = 20%. The red bars indicate the THF_30_ of a homogenous liver composed of functionally identical LUs.

### Patterning of steatosis as predictor of prospective NAFLD development

The heterogeneous distributions of intra-hepatic functional capacities result in different patterns of the regional TAG distribution during NAFLD progression. This raises the question whether the pattern of regional steatosis in an earlier phase of NAFLD progression may allow a prognosis of prospective disease development. We addressed this question by monitoring the TAG distribution across the individual LUs at 5, 10 and 20 years after onset of NAFLD. Livers with a severe outcome (TFH_30_ values lower than 55%) had on the average a strikingly higher proportion of non-steatotic LUs but the steatotic LUs had a higher mean TAG content than those in livers with a modest NAFLD outcome (TFH_30_ values larger than 75%). These two features of TAG distribution discriminating between severe and modest outcome of NAFLD can be captured by a *steatosis pattern score* (SPS),

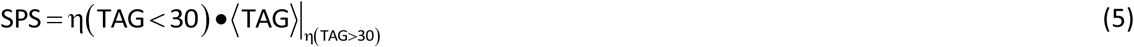

where the first factor represents the fraction of LUs with a TAG content lower than 30 mM and the second factor is the average TAG content of the complementary fraction of LUs with TAG content larger than 30 mM. Plotting TFH_30_ against the SPS (Fig. 6A-C) yielded significant negative correlations with increasing duration of the disease. For example, all livers with SPS < 16 at ten years after onset of NAFLD had THF_30_ values larger than 60% whereas all livers with SPS > 21 had THF_30_ values smaller than 60%. Thus, the regional steatosis pattern at earlier time points of NAFLD progression appears to contain valuable information of the further progression of the disease. This conclusion holds without saying for conditions where the FFA challenge persists and other disease-promoting hits are absent.

**Fig 6.**
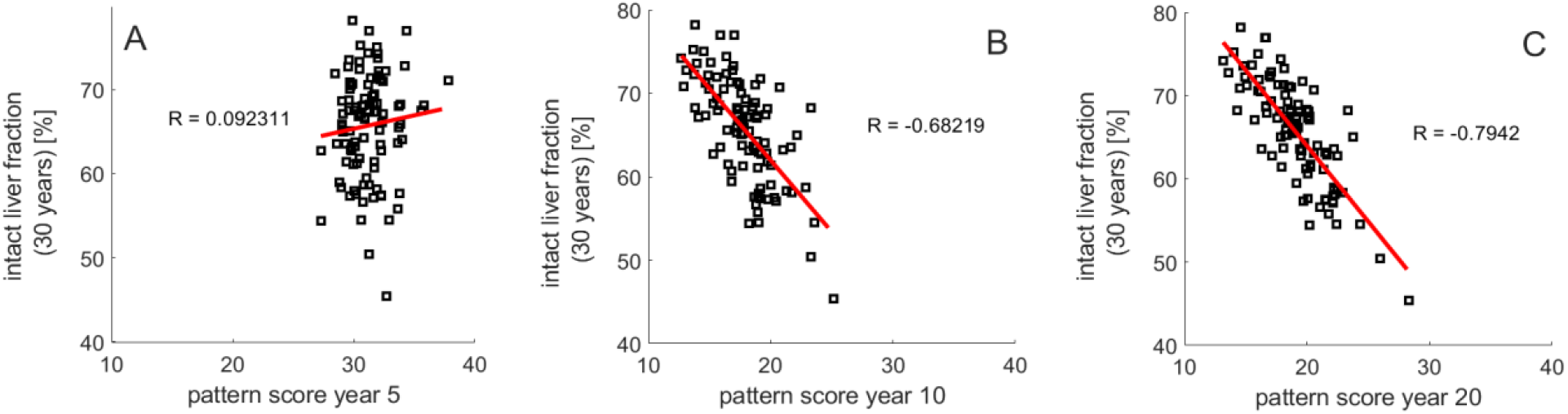
Relation between steatosis patterning and NAFLD progression. THF_30_ versus steatosis pattern score (SPS) defined by equation (5) at (A) 5 years, (B) 10 years and (C) 20 years. 100 simulations with the same parameters as in Fig. 3.

### Exacerbation of NAFLD by additional liver-damaging hits

In the preceding sections, simulated progression of NAFLD was exclusively driven by a FFA challenge without presence of further liver-damaging ‘hits’. Such hits may be caused, for example, by changes of the gut microbiome, alcohol consumption, xenobiotica, drugs or viral infections. As proposed by the ‘second hit’ hypothesis [51, 52] and more recently by the ‘multiple hit’ hypothesis [11], additional threats are necessary to drive an initially steatotic liver into NASH and even more severe stages of liver damage.

However, as suggested by the simulations shown in the preceding sections, for livers with an unfavorable intra-hepatic distribution of metabolic and repair capacities, such additional hits are not required to drive the liver into severe pathological states (see example in Fig. 4B). Nevertheless, occurrence of additional hits may result in a dramatic acceleration of NAFLD progression. This is illustrated in Fig. 7 for a liver that was exposed to three consecutive damaging hits with a duration of about one year. In the absence of the FFA challenge, the hits resulted in a decrease of the total hepatocyte fraction to about 85%, which was reversible because the timely distance between the hits (∼3 years in this example) was long enough to allow for a complete recovery. The FFA challenge of the same liver resulted in a partially damaged liver with a stable TFH_30_ of about 63%. If the fatty liver was additionally exposed to the same hits as the healthy liver, TFH dropped gradually to eventually fall below 30%. Of note, the extent of the hit-induced drop of the hepatocyte fraction became smaller with increasing number of hits, a feature that was consistently observed in these simulations. Every hit removes a fraction of LUs with less favorable damage-counteracting capacities so that, on the average, the damage resistance and repair capacity of the surviving LUs was increasing.

**Fig 7.**
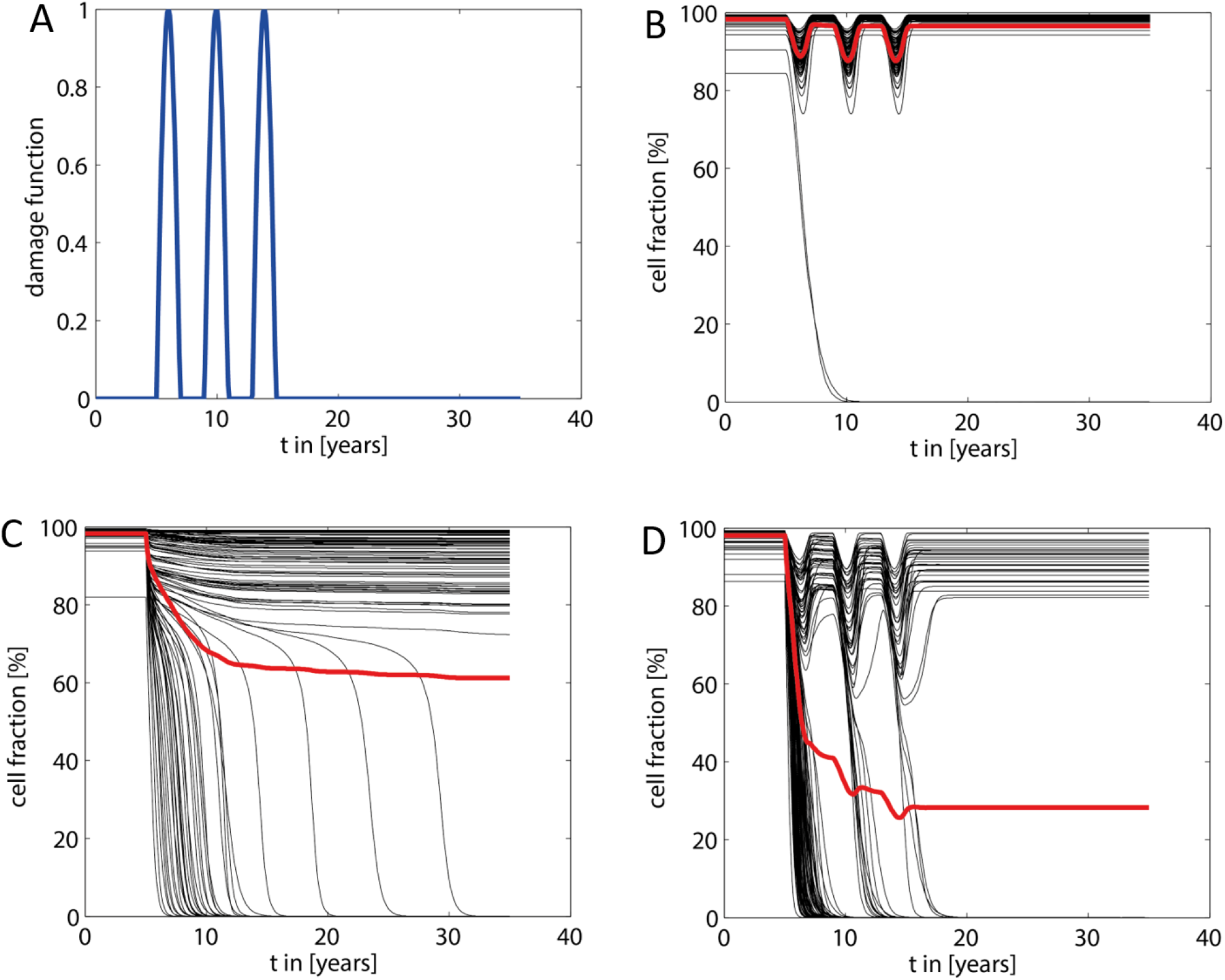
Acceleration of NAFLD progression by additional damaging ‘hits’. A) The additional ‘hits’ were modeled by a damage function H(t) consisting of three consecutive hits in a timely distance of three years, each hit having a duration of about one year and a maximal strength of 2.1·10^−3^ d^-1^, which is six times the basal damage rate k_d_. At each time point, H(t) was added to the damage function FD(t) defined in equation (2). B) Response of the healthy liver (no NAFLD) to the transient hits shown in A). C) Response of the liver to an increased FFA load (parameters the same as in Fig. 4). D) Response of the NAFLD liver shown in C) to the transient hits shown in A).

## Discussion

### Intra-hepatic distribution of functional capacities influences NAFLD progression

The most important finding of our computational study is that the heterogeneous intra-hepatic distribution of metabolic and tissue-remodeling capacities is a strong determinant of NAFLD progression. Whether the onset of fatty-acid induced cellular damage in a fraction of liver regions may trigger a cascading damage depends on the functional capacities of the unaffected regions to effectively handle FFAs. We hypothesize that an unfavorable intra-hepatic distribution of functional capacities can be sufficient to drive simple steatosis to severe forms of NAFLD without occurrence of additional hits. However, as with all diseases, numerous additional risk factors can accelerate worsening of the disease. Our simulations suggest that even in livers with mild NAFLD, the impact of such additional ‘hits’ is much more severe than in the healthy liver. Interestingly, the repeated occurrence of transient liver-damaging events represents a kind of selection pressure that favors the ‘survival’ of LUs possessing the largest capacities to withdraw and repair tissue damage (see Fig. 7D).

### Cascading failure as a fundamental principle of disease progression

Our study of NAFLD progression rests on the principle of cascading organ failure: Loss of function due to the failure of parts of an organ/tissue has to be compensated by its functionally intact parts. This in turn exerts an additional load to the intact parts thus causing them to fail as well if adaptive mechanisms have reached their limit. In our model, the adaptive response was implicitly taken into account by the a priori statistical distribution of functional parameters across the LUs. It is reasonable to assume that the principle of cascading organ failure applies to the disease progression in other organs, e.g. heart failure due to long-lasting hemodynamic overload, liver or kidney failure caused by drug intoxication or insulin resistance and inflammation of adipose tissue caused by a TAG load exceeding the expansion limit.

### Macro-scale versus micro-scale heterogeneity

Our mathematical model rests on the assumption that distinct spatial regions (that we named liver units – LUs) in the size range of centimeters exist, which differ from each other by blood perfusion rates, metabolic capacities of hepatocytes and the endowment with other cell types, in particular Kuppfer cells and fibroblasts involved in defense and repair processes. This macro-scale heterogeneity has to be distinguished from the well-studied micro-scale heterogeneity among different zones of the acinus, commonly referred to as metabolic (functional) zonation. The existence of functional macro-scale heterogeneity of the liver can be inferred from the spatially variable distribution of blood flow rates, lipid accumulation, and clearance rates of drugs and metabolites revealed by various imaging techniques. The molecular and cellular basis of these variations remains elusive. The vasculature of the liver may play an important role. For example, the pancreatic hormone insulin, which stimulates lipogenesis, has quite different concentrations in the various tributaries of the portal blood. Hence, hepatic territories low in insulin may explain areas spared by fat in a steatotic patient [53]. Concerning the contribution of regional differences in gene expression to intrahepatic functional heterogeneity, existing studies do not go beyond the single acinus although the technical prerequisites are now available [54]. Generally, it is well established that environmental fluctuations (extrinsic noise) affect the development of individual organisms and tissues [55]. Gene expression studies across mammalian organs suggest that random external events may largely influence the temporal trajectories of gene expression during organ development [56].

### Spatial patterning of steatosis: An additional risk factor?

Our model simulations suggest an association between the pattern of steatosis and the long-term outcome of NAFLD. A strongly contrasting steatosis patterning, i.e. the coexistence of non-steatotic and highly-steatotic regions, appears to be indicative for the later occurrence of severe NAFLD stages (see Fig. 6). In the light of this finding, it could be worth to carry out a retrospective study relating clinical and histopathological data on NAFLD progression to the spatial pattern of steatosis revealed by ultrasound or nuclear magnetic resonance at the time point of diagnosis. If such study supports the model-derived hypothesis, spatially-resolved patterning of steatosis by means of non-invasive imaging techniques might be included in the list of markers for the risk of a simple steatosis to progress to more severe stages of NAFLD.

### Further extensions of the model

The presented mathematical model of NAFLD progression is a generic one in that fundamental processes involved in the initiation and progression of the disease have been included in the model as events that trigger each other. Gradually including the huge variety of intertwined molecular and cellular processes underlying these events may seem as useful extension of the model. However, this extension should not be carried out according to the motto “put in everything that is known” but with a clear strategy that is oriented at a defined medical goal. One of the central medical goals in NAFLD research is to better understand and prevent the transition from simple steatosis to NASH. In this respect, the more detailed modeling of mechanisms and pathways involved in lipotoxic cell damage appears to be crucial in view of controversial ideas on useful pharmacological interventions. For example, abrogating the increase of hepatic lipogenesis (e.g. by inactivation of SREBP) has been advocated as a promising way to prevent steatosis and thus NASH [5]. According to our simulations (see Fig. 2) and experimental findings [4], this might be the wrong way because enhanced lipogenesis is an important defense mechanism to lower the concentration of potentially toxic FFAs. On the other hand, excessive accumulation of TAG may induce endoplasmic reticulum stress and cell damage [57]. This Janus-faced effect of hepatic TAG synthesis on the progression of NAFLD calls for a more detailed and physiology-based model of the cellular lipid metabolism, which in particular includes the various processes involved in the synthesis, growth and degradation of lipid droplets [58]. Another important aspect of NAFLD progression, which was neglected in our model, is the upregulation of metabolic pathways to compensate for the decline of active hepatocyte mass. For example, It is well established that pro-inflammatory cytokines have a strong impact of on the lipid metabolism of the liver [59]. In the inflammatory phase of the disease, cytokine-stimulated upregulation of lipogenic pathways could facilitate the incorporation of potentially toxic free FAs into TAG.

## Author Contributions

Conceptualization, HGH; Methodology, HGH and NB; Software, NB; Validation, HGH and NB; Formal Analysis, HGH and NB; Investigation, NB; Resources, HGH; Data Curation, NB; Writing – Original Draft Preparation, HGH; Writing – Review & Editing, HGH and NB; Visualization, HGH and NB; Project Administration, HGH; Funding Acquisition, HGH.

## References

1. DeWeerdt, S., Disease progression: Divergent paths. Nature, 2017. 551(7681).

2. Sondergaard, E. and M.D. Jensen, Quantification of adipose tissue insulin sensitivity. J Investig Med, 2016. 64(5): p. 989–91.

3. Choi, S.S. and A.M. Diehl, Hepatic triglyceride synthesis and nonalcoholic fatty liver disease. Curr Opin Lipidol, 2008. 19(3): p. 295–300.

4. Listenberger, L.L., et al., Triglyceride accumulation protects against fatty acid-induced lipotoxicity. Proc Natl Acad Sci U S A, 2003. 100(6): p. 3077–82.

5. Papazyan, R., et al., Physiological Suppression of Lipotoxic Liver Damage by Complementary Actions of HDAC3 and SCAP/SREBP. Cell Metab, 2016. 24(6): p. 863–874.

6. Yamaguchi, K., et al., Inhibiting triglyceride synthesis improves hepatic steatosis but exacerbates liver damage and fibrosis in obese mice with nonalcoholic steatohepatitis. Hepatology, 2007. 45(6): p. 1366–74.

7. Liu, J., et al., Free fatty acids, not triglycerides, are associated with non-alcoholic liver injury progression in high fat diet induced obese rats. Lipids Health Dis, 2016. 15: p. 27.

8. Ibrahim, S.H., R. Kohli, and G.J. Gores, Mechanisms of lipotoxicity in NAFLD and clinical implications. J Pediatr Gastroenterol Nutr, 2011. 53(2): p. 131–40.

9. Cordero-Espinoza, L. and M. Huch, The balancing act of the liver: tissue regeneration versus fibrosis. J Clin Invest, 2018. 128(1): p. 85–96.

10. Seko, Y., K. Yamaguchi, and Y. Itoh, The genetic backgrounds in nonalcoholic fatty liver disease. Clin J Gastroenterol, 2018. 11(2): p. 97–102.

11. Tilg, H. and A.R. Moschen, Evolution of inflammation in nonalcoholic fatty liver disease: the multiple parallel hits hypothesis. Hepatology, 2010. 52(5): p. 1836–46.

12. Berndt, N., et al., Functional Consequences of Metabolic Zonation in Murine Livers: Insights for an Old Story. Hepatology, 2020.

13. Halpern, K.B., et al., Erratum: Single-cell spatial reconstruction reveals global division of labour in the mammalian liver. Nature, 2017. 543(7647): p. 742.

14. Malarkey, D.E., et al., New insights into functional aspects of liver morphology. Toxicol Pathol, 2005. 33(1): p. 27–34.

15. Berndt, N. and H.G. Holzhutter, Dynamic Metabolic Zonation of the Hepatic Glucose Metabolism Is Accomplished by Sinusoidal Plasma Gradients of Nutrients and Hormones. Front Physiol, 2018. 9: p. 1786.

16. Gebhardt, R. and M. Matz-Soja, Liver zonation: Novel aspects of its regulation and its impact on homeostasis. World J Gastroenterol, 2014. 20(26): p. 8491–504.

17. Kleiner, D.E. and H.R. Makhlouf, Histology of Nonalcoholic Fatty Liver Disease and Nonalcoholic Steatohepatitis in Adults and Children. Clin Liver Dis, 2016. 20(2): p. 293–312.

18. Sherriff, S.B., R.C. Smart, and I. Taylor, Clinical study of liver blood flow in man measured by 133Xe clearance after portal vein injection. Gut, 1977. 18(12): p. 1027–31.

19. Wang, H., et al., Predictive models for regional hepatic function based on 99mTc-IDA SPECT and local radiation dose for physiologic adaptive radiation therapy. Int J Radiat Oncol Biol Phys, 2013. 86(5): p. 1000–6.

20. Sorensen, M., et al., Regional metabolic liver function measured in patients with cirrhosis by 2-[(1)(8)F]fluoro-2-deoxy-D-galactose PET/CT. J Hepatol, 2013. 58(6): p. 1119–24.

21. Bonekamp, S., et al., Spatial distribution of MRI-Determined hepatic proton density fat fraction in adults with nonalcoholic fatty liver disease. J Magn Reson Imaging, 2014. 39(6): p. 1525–32.

22. Hamer, O.W., et al., Fatty liver: imaging patterns and pitfalls. Radiographics, 2006. 26(6): p. 1637–53.

23. Jensen, V.S., et al., Variation in diagnostic NAFLD/NASH read-outs in paired liver samples from rodent models. J Pharmacol Toxicol Methods, 2020. 101: p. 106651.

24. Berndt, N., et al., A multiscale modelling approach to assess the impact of metabolic zonation and microperfusion on the hepatic carbohydrate metabolism. PLoS Comput Biol, 2018. 14(2): p. e1006005.

25. Soler-Argilaga, C. and M. Heimberg, Comparison of metabolism of free fatty acid by isolated perfused livers from male and female rats. J Lipid Res, 1976. 17(6): p. 605–15.

26. Viljanen, A.P., et al., Effect of weight loss on liver free fatty acid uptake and hepatic insulin resistance. J Clin Endocrinol Metab, 2009. 94(1): p. 50–5.

27. Weisiger, R., J. Gollan, and R. Ockner, Receptor for albumin on the liver cell surface may mediate uptake of fatty acids and other albumin-bound substances. Science, 1981. 211(4486): p. 1048–51.

28. Lambert, J.E., et al., Increased de novo lipogenesis is a distinct characteristic of individuals with nonalcoholic fatty liver disease. Gastroenterology, 2014. 146(3): p. 726–35.

29. Iozzo, P., et al., Liver uptake of free fatty acids in vivo in humans as determined with 14(R, S)-[18F]fluoro-6-thia-heptadecanoic acid and PET. Eur J Nucl Med Mol Imaging, 2003. 30(8): p. 1160–4.

30. Vatner, D.F., et al., Insulin-independent regulation of hepatic triglyceride synthesis by fatty acids. Proc Natl Acad Sci U S A, 2015. 112(4): p. 1143–8.

31. Soler-Argilaga, C. and M. Heimberg, Calculation of the rate of utilization of albumin-bound free fatty acids from specific radioactivity data. Lipids, 1976. 11(1): p. 82–4.

32. Mittendorfer, B., et al., VLDL Triglyceride Kinetics in Lean, Overweight, and Obese Men and Women. J Clin Endocrinol Metab, 2016. 101(11): p. 4151–4160.

33. Goh, E.H. and M. Heimberg, Effects of free fatty acids on activity of hepatic microsomal 3-hydroxy-3-methylglutaryl coenzyme A reductase and on secretion of triglyceride and cholesterol by liver. J Biol Chem, 1977. 252(9): p. 2822–6.

34. Ipsen, D.H., J. Lykkesfeldt, and P. Tveden-Nyborg, Molecular mechanisms of hepatic lipid accumulation in non-alcoholic fatty liver disease. Cell Mol Life Sci, 2018. 75(18): p. 3313–3327.

35. Schlierf, G., W. Reinheimer, and V. Stossberg, Diurnal patterns of plasma triglycerides and free fatty acids in normal subjects and in patients with endogenous (type IV) hyperlipoproteinemia. Nutr Metab, 1971. 13(2): p. 80–91.

36. Quinn, W.J., 3rd, et al., mTORC1 stimulates phosphatidylcholine synthesis to promote triglyceride secretion. J Clin Invest, 2017. 127(11): p. 4207–4215.

37. Shimoda, T., et al., A low coefficient of variation in hepatic triglyceride concentration in an inbred rat strain. Lipids Health Dis, 2020. 19(1): p. 137.

38. Malhi, H. and G.J. Gores, Molecular mechanisms of lipotoxicity in nonalcoholic fatty liver disease. Semin Liver Dis, 2008. 28(4): p. 360–9.

39. Mendez-Sanchez, N., et al., New Aspects of Lipotoxicity in Nonalcoholic Steatohepatitis. Int J Mol Sci, 2018. 19(7).

40. Macdonald, R.A., “Lifespan” of liver cells. Autoradio-graphic study using tritiated thymidine in normal, cirrhotic, and partially hepatectomized rats. Arch Intern Med, 1961. 107: p. 335–43.

41. Burczynski, F.J., B.A. Luxon, and R.A. Weisiger, Intrahepatic blood flow distribution in the perfused rat liver: effect of hepatic artery perfusion. Am J Physiol, 1996. 271(4 Pt 1): p. G561–7.

42. Zhang, X., et al., Proteomic analysis of individual variation in normal livers of human beings using difference gel electrophoresis. Proteomics, 2006. 6(19): p. 5260–8.

43. Zhu, Y., et al., Identification of Protein Abundance Changes in Hepatocellular Carcinoma Tissues Using PCT-SWATH. Proteomics Clin Appl, 2019. 13(1): p. e1700179.

44. Puri, P., et al., A lipidomic analysis of nonalcoholic fatty liver disease. Hepatology, 2007. 46(4): p. 1081–90.

45. Meyerson, C. and B.V. Naini, Something old, something new: liver injury associated with total parenteral nutrition therapy and immune checkpoint inhibitors. Hum Pathol, 2020. 96: p. 39–47.

46. Yki-Jarvinen, H., et al., Severity, duration, and mechanisms of insulin resistance during acute infections. J Clin Endocrinol Metab, 1989. 69(2): p. 317–23.

47. Lytle, K.A. and D.B. Jump, Is western diet-induced nonalcoholic steatohepatitis in Ldlr-/-mice reversible? PloS one, 2016. 11(1): p. e0146942.

48. Ekstedt, M., et al., Long-term follow-up of patients with NAFLD and elevated liver enzymes. Hepatology, 2006. 44(4): p. 865–73.

49. Bashir, M.R., et al., Quantification of hepatic steatosis with a multistep adaptive fitting MRI approach: prospective validation against MR spectroscopy. AJR Am J Roentgenol, 2015. 204(2): p. 297–306.

50. Reeder, S.B., et al., Quantification of hepatic steatosis with MRI: the effects of accurate fat spectral modeling. J Magn Reson Imaging, 2009. 29(6): p. 1332–9.

51. Basaranoglu, M., G. Basaranoglu, and H. Senturk, From fatty liver to fibrosis: a tale of “second hit”. World J Gastroenterol, 2013. 19(8): p. 1158–65.

52. Mole, D.J., et al., The isolated perfused liver response to a ‘second hit’ of portal endotoxin during severe acute pancreatitis. Pancreatology, 2005. 5(4-5): p. 475–85.

53. Vilgrain, V., et al., Hepatic steatosis: a major trap in liver imaging. Diagn Interv Imaging, 2013. 94(7-8): p. 713–27.

54. Zhu, Q., et al., Identification of spatially associated subpopulations by combining scRNAseq and sequential fluorescence in situ hybridization data. Nat Biotechnol, 2018.

55. Tsimring, L.S., Noise in biology. Rep Prog Phys, 2014. 77(2): p. 026601.

56. Cardoso-Moreira, M., et al., Gene expression across mammalian organ development. Nature, 2019. 571(7766): p. 505–509.

57. Han, J. and R.J. Kaufman, The role of ER stress in lipid metabolism and lipotoxicity. J Lipid Res, 2016. 57(8): p. 1329–38.

58. Wallstab, C., et al., A unifying mathematical model of lipid droplet metabolism reveals key molecular players in the development of hepatic steatosis. FEBS J, 2017. 284(19): p. 3245–3261.

59. Hijona, E., et al., Inflammatory mediators of hepatic steatosis. Mediators Inflamm, 2010. 2010: p. 837419.

